# A supervised digital game intervention supports language and communication in young children

**DOI:** 10.64898/2026.04.02.716239

**Authors:** Marcela Peña, Ghislaine Dehaene-Lambertz, Esteban J. Pino, Enrica Pittaluga, Patricia Cortés, Consuelo de la Riva, Orieta Palacios, Pamela Guevara

## Abstract

The role of digital media in early childhood development remains highly debated, particularly regarding its impact on language acquisition. While excessive or unsupervised screen exposure has been linked to poorer outcomes, less is known about whether structured and interactive uses of technology can support learning. Building on previous research, we evaluated a brief, educator-supervised tablet-based intervention in 246 children aged 2-5 years from low- to middle-socioeconomic backgrounds attending public early education centers. Using a pre-post design with matched study and control groups, children completed 4-8 short training sessions (15 minutes each) involving interactive word-image associations spanning multiple linguistic categories. Preschoolers additionally engaged in prompted vocalization.

Across age groups (2-3, 3-4, and 4-5 years), children in the intervention showed greater gains in language comprehension than controls, including receptive language in toddlers (β = 0.49, p = 0.009), vocabulary and morphology in younger preschoolers (β = 0.59-0.68, all p < 0.05), and grammar comprehension in older preschoolers (β = 0.30, p = 0.038). These effects were consistent after accounting for child and parental characteristics.

Together, these findings suggest that the developmental impact of digital media depends less on exposure itself than on how it is used. When embedded in structured, socially guided interactions, even brief tablet-based activities may support early language development

## Introduction

Despite the widespread use of interactive technologies by young children, clear guidelines on how to maximize their educational benefits while minimizing potential harm remain limited [1]. Concerns about early screen exposure have been widely discussed, particularly in relation to language development and broader neurodevelopmental outcomes. While some studies report associations between high levels of screen time and adverse developmental outcomes, including language delays [2,3], the evidence remains mixed, and causal relationships are difficult to establish due to confounding environmental and familial factors [4,5].

A key distinction emerging from literature is between passive and interactive forms of screen use. Passive exposure, particularly when it displaces caregiver-child interaction, has been associated with reduced verbal engagement and poorer language outcomes in early childhood [2]. In contrast, interactive and socially scaffolded uses of digital media, especially those involving contingent feedback or adult participation, appear to mitigate these risks and may even support aspects of cognitive and language development [6,7].

From a developmental perspective, early language acquisition is highly sensitive to the quality of environmental input, including the richness of linguistic interactions and opportunities for contingent learning [8,9]. Digital tools that incorporate features such as immediate feedback, multimodal input, and guided attention may therefore provide structured opportunities to support language learning, particularly in contexts where access to enriched linguistic environments is limited. This is especially relevant in low-to middle-income settings, where scalable and cost-effective interventions are needed.

Building on previous work demonstrating that brief, tablet-based training can support early communicative abilities [10], the present study extends this approach by targeting a broader range of linguistic structures, including verbs, adjectives, adverbs, and passive constructions. By more closely approximating the complexity of natural language, this expanded intervention aims to assess whether structured, supervised digital interactions can be effectively leveraged to support early language development.

The primary objective of this study was to evaluate whether a short, educator-supervised tablet-based intervention is associated with improvements in language comprehension in toddlers and preschoolers. Given the ongoing debate surrounding digital media use in early childhood, this work seeks to contribute empirical evidence regarding the conditions under which such technologies may support, rather than hinder, developmental outcomes. This perspective emphasizes that the developmental impact of digital media depends less on exposure per se than on the quality and context of its use.

## Materials and Methods

### Study Design

We implemented a quasi-experiment design, consisting of a training period with pre- and post-training evaluations of linguistic abilities. The study included toddlers (aged 2-3 years) and preschoolers with the latter divided into two distinct cohorts: 3-to-4-year-olds and 4-to-5-year-olds. These three groups (thereafter referred to by their respective age ranges) were analyzed separately to account for developmental differences.

We assigned children to the study (hereafter Study) or Control group, balancing by pre-training scores language scores, child’s sex and age and mother’s age and education.

Because for ethical reasons a training phase was planned for the Control group, to take place after the end of the post-training step, sanitary restrictions prevented its implementation with consistent frequency and conditions (the winter break in preschools was advanced 3 weeks). We remark that sanitary restrictions did not affect the comparison between study and control groups at the pre and post training steps. Our study was approved by the institutional review board of the Pontificia Universidad Católica de Chile. Parents provided written informed consent before their child participated.

### Participants

Children were recruited from eight early childhood care centers in Santiago, Chile, between 2021 and 2023, a period characterized by pandemic-related health restrictions. All participants came from low-to middle-socioeconomic-status, monolingual Spanish-speaking families. Children were born full-term, exhibited typical developmental trajectories at the time of intervention, and had no reported family history of language impairment or dyslexia.

Exclusion criteria included diagnosed visual or auditory impairments, neurodevelopmental disorders, and chronic conditions associated with developmental delays. Children with neonatal asphyxia, epilepsy, chromosomal abnormalities, or inborn errors of metabolism were excluded.

Of the 565 families invited, 332 provided consent to participate. The final analytical sample consisted of 246 children who completed pre- and post-training evaluations for at least one assessment battery. For those in the study groups, inclusion required participation in four to eight training sessions. The final cohort comprised 87 toddlers (2-3 years), 85 preschoolers aged 3-4 years, and 74 aged 4-5 years. Participant dropout was largely due to public health restrictions related to seasonal respiratory illnesses including SARS-CoV-2. The excluded and included children did not differ in demographics characteristics (Supporting information Table S1).

### Measures

Pre-post training evaluations included parental reports and direct child assessments using age-appropriate standardized batteries.

Parental reports included the Spanish versions of the Ages & Stages Questionnaires [11,12] (hereafter ASQ-3), which evaluates Communication, Gross Motor, Fine Motor, Problem Solving, and Personal-Social domains. ASQ-3 data was collected for all participants.

For 2-to-3-years, we also administered the short form of the MacArthur-Bates Communicative Development Inventories [13,14] (hereafter CDI), a parent-report questionnaire assessing expressive vocabulary (via a 100-word checklist) and early speech complexity (use of two-words combinations). Additionally, a child psychologist administered the Receptive and Expressive language scales of the Bayley Scales of Infant and Toddler Development, 3^rd^ Edition, to each child.

For 3-to-4- and 4-to-5-years, a speech therapist administered the Screening Test of Spanish Grammar [15] (hereafter, STSG) to evaluate receptive and expressive grammar.

Additionally, 3-to-4-years completed the Spanish version of the Test for Auditory Comprehension of Language [16] (hereafter TACL), chosen for its age-appropriate, concise instructions regarding vocabulary, morphology, and syntax. 4-to-5-years completed the Phonological Awareness Assessment [17] (hereafter PAA) on an exploratory basis to estimate emergent pre-reading skills, specifically syllabic and phonemic awareness.

Except for CDI, all assessment tools provide both raw scores and age-normed categorical classifications. Since most participants fell within normative limits, we analyzed raw scores to capture finer variations in performance. Child psychologists and speech therapists were external to the research team and remained blind to the group assignation of each child.

### Procedures

The procedure was similar to the one reported in (ref. 10). Pre- and post-training evaluations involved parent-completed questionnaires and professional assessments administered by child psychologists (toddlers) and speech therapists (preschoolers). To minimize bias, all evaluators were external to the study and blinded to experimental conditions.

The training was administered individually and supervised by an early childhood educator in a quiet room. During the training, a series of videos featuring an early childhood educator guided the children in associating spoken words with their corresponding images (Fig. 1). Contrary to the previous study (ref. 8), these associations spanned a diverse linguistic range, including nouns, verbs, adjectives, and adverbs of place, manner, or quantity, as well as passive constructions. The training consisted of four to eight 15-minute sessions, each including five trials. Each trial was divided in two phases: encoding and recognition. During encoding, children established associations between two spoken words and their corresponding images through interactive touch-screen prompts. During recognition, they were asked to match one of the trained words to one of the two images presented simultaneously. Tactile responses were recorded and classified as correct (target selection), omitted (no response), or incorrect (selection of the distractor or an area outside the target boundaries). The training was guided by videos of an early childhood educator who invited the child to play using direct child speech, direct gaze and smiling gestures during every trial and provided immediate encouraging social feedback after each response, regardless of the correctness.

**Figure 1.**
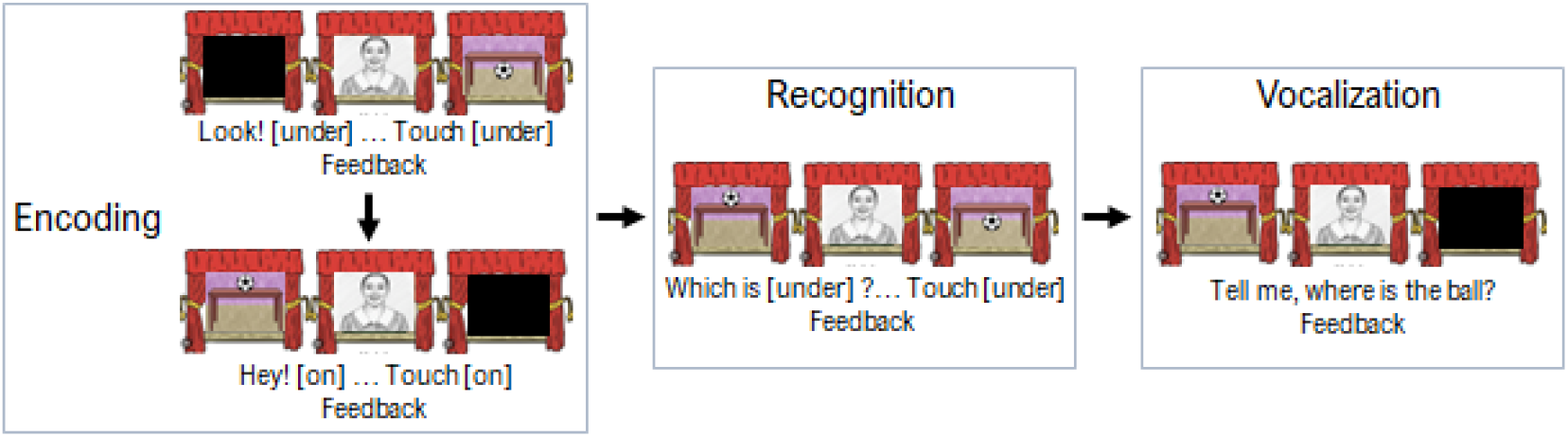
Schematic structure of a training trial. The training was guided by a prerecorded video of an early childhood educator providing child-directed instructions and feedback. Each trial comprised three phases. During the encoding phase, children were presented with word-image pairs and prompted to select the corresponding image via touchscreen interaction. In the recognition phase, children were asked to identify previously presented associations. For children older than 3 years, a vocalization phase was included, in which the educator prompted the child to produce a spoken response. All phases included immediate, non-corrective feedback to maintain engagement.

In the preschool cohorts, trials concluded with a vocalization phase wherein the educator in the video prompted the child to produce the target word. To encourage engagement to speak, any speech-like vocalization was categorized as a correct response and replayed to the child as auditory reinforcement. Conversely, omitted or incorrect responses triggered encouraging, non-corrective feedback.

### Statistical Analysis

To assess the impact of the intervention, we computed standardized gain scores for each assessment battery within each age group. Standardized gain was defined as the difference between post- and pre-training raw scores divided by the pooled pre-training standard deviation of the corresponding age group, irrespective of group assignment. The mean interval between assessments was 59 days for toddlers and 60 days for preschoolers.

To account for non-normality in gain scores, we employed nonparametric rank-based methods. Specifically, we used rank-based linear regression implemented in the Rfit package in R [18] to evaluate the effect of the intervention while adjusting for potential confounding variables. Unlike ordinary least squares regression, this approach is robust to violations of normality and to the presence of outliers, as it minimizes Jaeckel’s dispersion function using Wilcoxon scores [19]. Given the developmental nature of our sample, p-values and standard errors for regression coefficients (β) were estimated using a permutation-based approach with 10,000 repetitions to ensure stable inference.

The final analysis included all children who provided both pre- and post-intervention data for at least one measure; participants in the intervention group were additionally required to complete a minimum of four training sessions. We also conducted an exploratory analysis of vocalizations produced by preschoolers in the intervention group.

Given the number of outcome measures and age-stratified analyses, the analyses were considered exploratory. Accordingly, we did not apply formal corrections for multiple comparisons and instead interpreted results with appropriate caution, emphasizing consistency across measures and effect sizes rather than isolated p-values.

## Results

At baseline, no significant differences were observed between groups regarding bio-demographic characteristics (Table 1). This equivalence ensures that the cohorts were comparable prior to the intervention.

**Table 1.**
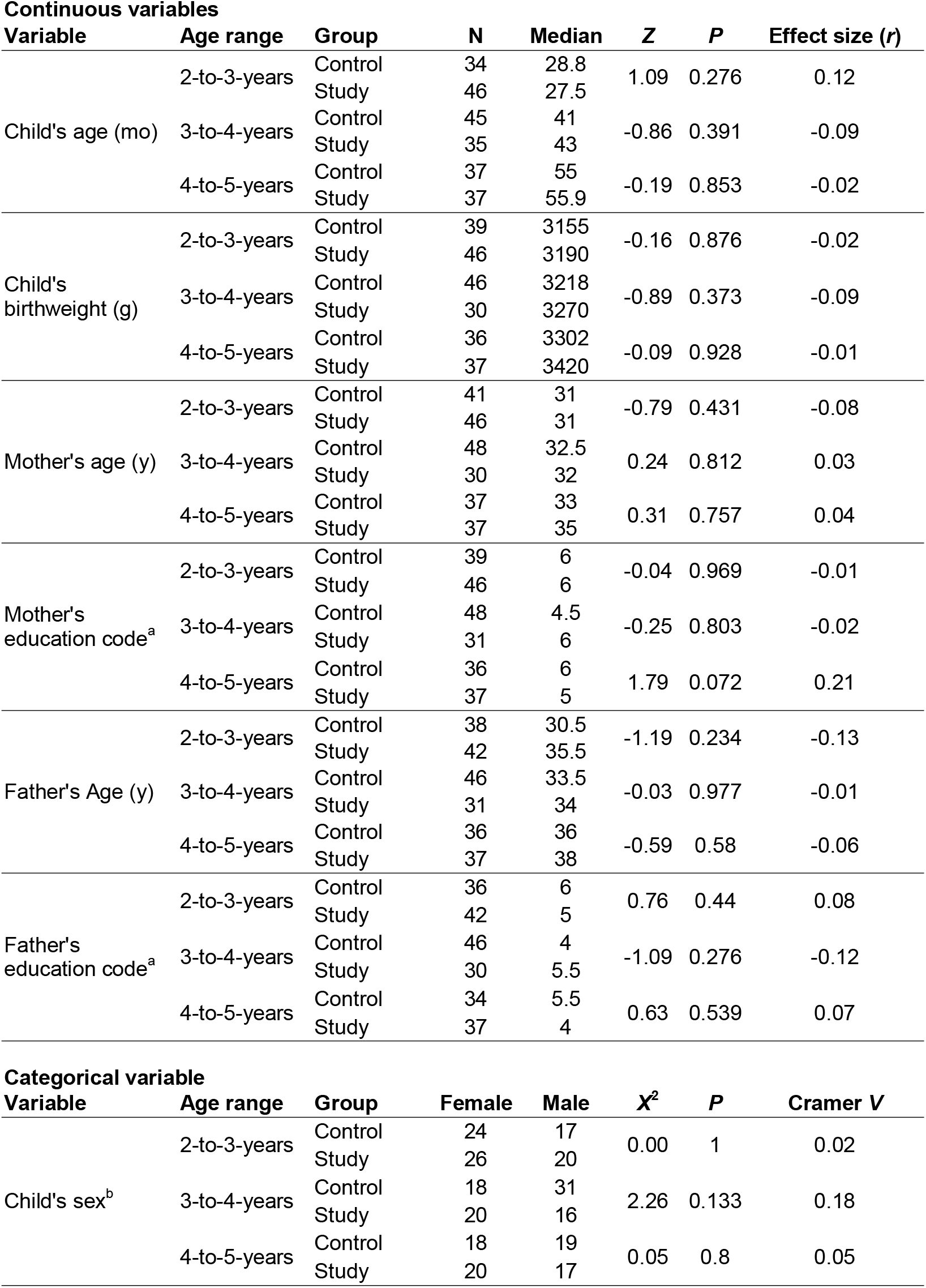
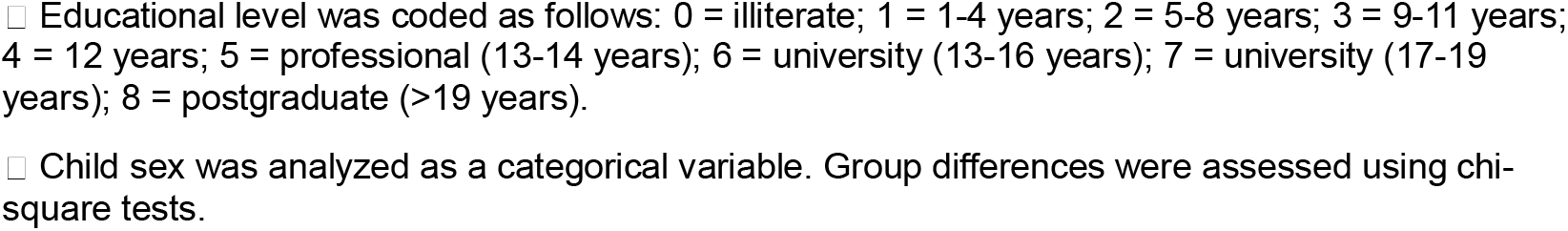
Baseline demographic characteristics by group at pre-training evaluation.

### Linguistic gains

#### 2-to-3-years

Analyses of the ASQ-3 (Fig. 2A) and CDI revealed no significant differences between groups, suggesting that parental perceptions of general developmental progress were similar across both groups.

**Figure 2.**
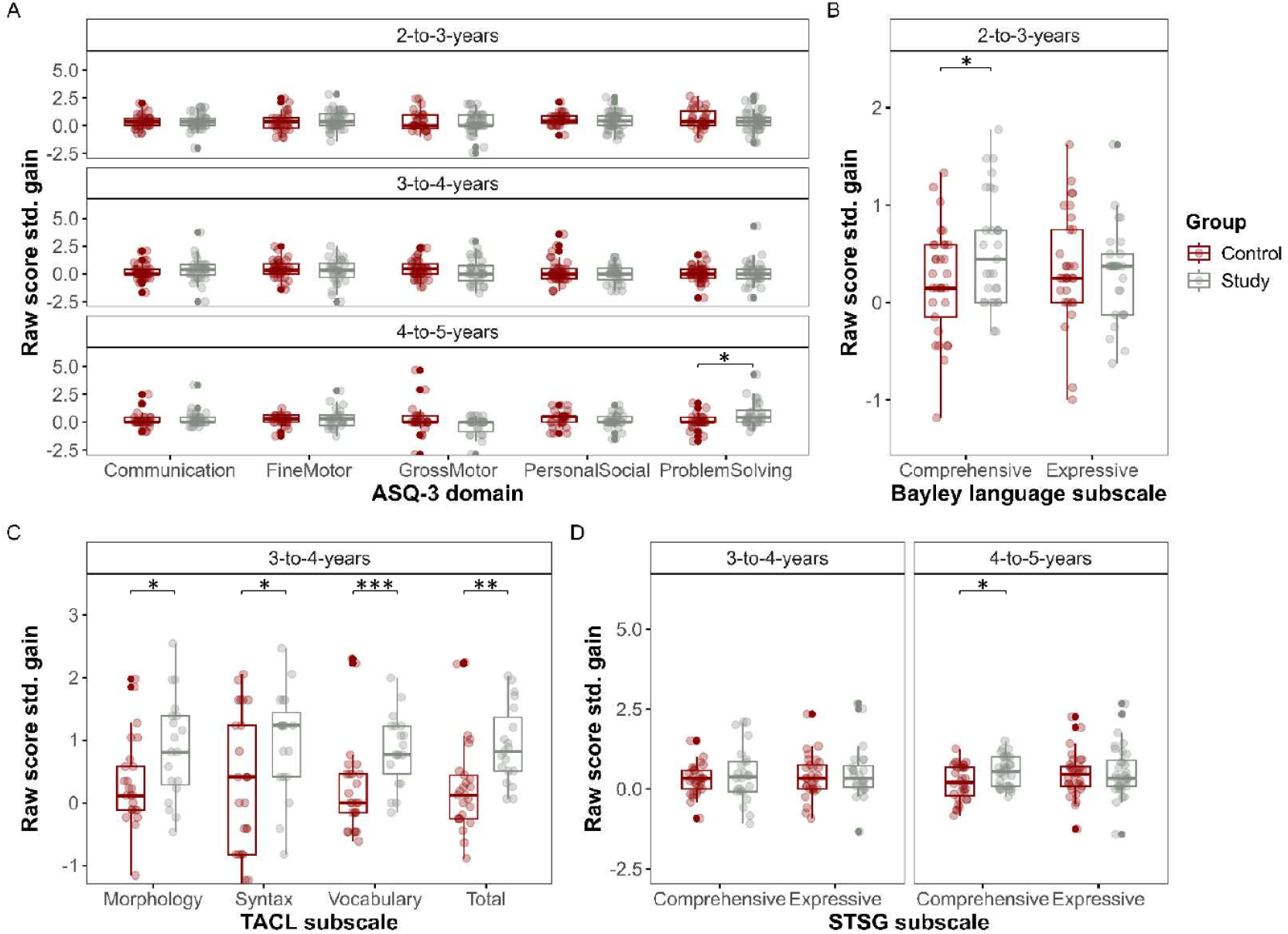
Developmental and language gains by age and domain. In **A**, we plot the standardized gains of raw scores in ASQ-3 domains for ages 2-3, 3-4, and 4-5 years. Control and Study groups are plotted in red and grey, respectively. Study group showed higher gain in Problem solving for 4-5 years. In **B** we depict the gain on Bayley Language subscales for 2-3-year-olds. Study group shows greater gains in Language comprehension. In **C** we show the gains on TACL subscales and total score for 3-4-year-olds. Study group showed higher gain across all measures. In **D** we plot the gains on STSG subscales for 3-4 and 4-5-year-olds. Study group showed greater gain in Comprehensive subscale for 4-5 years. Boxplots show medians, interquartile ranges, and individual scores. Significance code: *: *p* < 0.05, **: *p* < 0.01, ***: *p* < 0.001.

In contrast, unadjusted analyses of the direct child assessment with Bayley Receptive Language scale showed that the Study group achieved significantly higher gain scores than the Control group (Median 0.52 vs. 0.15, Wilcoxon-Mann-Whitney Z = -2.01, p = 0.044, r = -0.22; Fig. 2B).

To account for potential confounding bio-demographic factors, we conducted a rank-based regression adjusting for toddler’s age and sex, as well as parental age and education. The effect of Group remained robust after adjustment (β = 0.49, SE = 0.18, p = 0.009), whereas no other covariates reached significance (all p > 0.30) (Table 2, 2-to-3-years section), indicating that language gains were significantly associated with the intervention.

**Table 2.** Robust rank-based regression models for linguistic gains. *B* represents unstandardized regression coefficients, indicating the change in the outcome for each one-unit increase in the predictor. For vocabulary gain, child age is reported as a standardized coefficient (β) to allow direct comparison with the factor Group. *SE* denotes the standard error. Overall model fit was assessed using the reduction in dispersion test (*p*-value). Robust *R*^2^ indicates the proportion of dispersion explained by the rank-based model. Significance’s codes: *: *p* < 0 .05; **: *p* < 0.01.

**(A) Ages 2-3 years (Bayley Scales)**
| Predictor | $\beta$ | SE | *T* | *p*-value |
| --- | --- | --- | --- | --- |
| Intercept | 0.71 | 0.98 | 0.73 | 0.473 |
| Group (Study) | 0.49 | 0.18 | 2.74 | 0.009** |
| Child age | 0.02 | 0.03 | 0.75 | 0.46 |
| Gender (Male) | 0.01 | 0.18 | 0.05 | 0.957 |
| Mother's education | 0.01 | 0.06 | 0.08 | 0.934 |
| Mother's age | -0.03 | 0.03 | -1.04 | 0.306 |
| Father's education | -0.06 | 0.06 | -0.97 | 0.338 |
| Father's age | 0 | 0.02 | 0.16 | 0.872 |

**(B) Ages 3-4 years (TACL)**
| Predictor | Morphology ($\beta$) | *p*-value | Syntax ($\beta$) | *p*-value | Vocabulary ($\beta$) | *p*-value |
| --- | --- | --- | --- | --- | --- | --- |
| Intercept | -1.2 | 0.603 | -2.33 | 0.561 | -0.09 | 0.913 |
| Group (Study) | 0.68 | 0.014* | 0.59 | 0.195 | 0.59 | 0.018* |
| Child age | 0.01 | 0.779 | 0.04 | 0.604 | 0.24 | 0.046* |
| Gender (Male) | -0.32 | 0.273 | 0.19 | 0.701 | -0.23 | 0.387 |
| Mother's education | 0.07 | 0.384 | 0.03 | 0.8 | 0.06 | 0.443 |
| Mother's age | 0 | 0.934 | 0 | 0.943 | 0.03 | 0.408 |
| Father's education | 0.09 | 0.26 | 0.04 | 0.798 | 0.06 | 0.431 |
| Father's age | 0.01 | 0.845 | 0.02 | 0.706 | -0.03 | 0.319 |

**(C) Ages 4-5 years (STSG)**
| Predictor | $\beta$ | SE | *T* | *p*-value |
| --- | --- | --- | --- | --- |
| Intercept | -0.17 | 0.82 | -0.21 | 0.834 |
| Group (Study) | 0.3 | 0.14 | 2.13 | 0.038* |
| Child age | 0.01 | 0.01 | 0.64 | 0.526 |
| Gender (Male) | 0.04 | 0.15 | 0.29 | 0.774 |
| Mother's education (z) | 0.18 | 0.08 | 2.11 | 0.039* |
| Mother's age | -0.03 | 0.02 | -1.28 | 0.206 |
| Father's education (z) | -0.2 | 0.08 | -2.38 | 0.021* |
| Father's age | 0.02 | 0.02 | 1.07 | 0.292 |

*Model fit: Robust $R^2 = 0.19$ ; Dispersion test = 1.75, $p = 0.118$*

So far, these results suggest that the Study group showed robust improvements in receptive language abilities. While the intervention was associated with these gains, bio-demographic variables, specifically child age and parental education, exerted negligible influence in our data.

#### 3-to-4-years

The Study group showed greater gains across all TACL subscales compared to the Control group (Fig. 2C). Gains were significantly higher in Morphology (median = 0.81 vs. 0.12, Z = −2.29, p = 0.021, r = −0.25), Syntax (median = 1.23 vs. 0.41, Z = −1.97, p = 0.048, r = −0.21), and Vocabulary (median = 0.77 vs. 0, Z = −3.41, p < 0.001, r = −0.37). The total linguistic gain, reported for clinical comparison only, was also higher in the Study group (median = 0.82 vs. 0.13, Z = −3.19, p = 0.001, r = −0.35). These results indicate that the intervention had a moderate effect on overall language comprehension, with the largest effect observed in the Vocabulary subscale.

As in toddlers, rank-based regression confirmed that Group was the strongest predictor of gain, with significant effects for Morphology (B = 0.679; p = 0.014) and Vocabulary (β = 0.593; p = 0.018) (Table 2, 3-to-4-year-olds section). Child age contributed to Vocabulary gain (β = 0.242; p = 0.046), but the effect of Group was more than twice as large, suggesting that the intervention was the primary driver of improvements at this age.

No significant differences were found in gains on the ASQ-3 or STSG. The lack of differences in the STSG is likely due to the longer and more complex instructions, which may be difficult for younger preschoolers to comprehend.

Together, these results indicate that 3-to-4-year-olds benefited from the intervention in language comprehension, with child age playing a secondary role.

#### 4-to-5-years

Children in the Study group showed significantly greater gains in language comprehension compared to those in the Control group (median = 0.55 vs. 0.21, respectively; Z = −2.14, p = 0.032, r = −0.25) (Fig. 2D). This effect remained significant after adjusting for child and parental bio-demographic factors (B = 0.30, SE = 0.14; t = 2.13; p = 0.038) (Table 2, 4-5-year-olds section).

The Study group also showed higher gains in the ASQ-3 Problem Solving domain (median = 0.43 vs. 0; Z = −2.29; p = 0.021; r = −0.27), but this effect became non-significant after adjusting for covariates (β = 0.40, SE = 0.23; t = 1.72; p = 0.092), suggesting that the intervention had only a marginal effect on this domain.

No other significant differences were observed (all p > 0.05).

Importantly, across age groups, the intervention was not associated with negative effects. Instead, all Study groups showed small-to-moderate but consistent improvements in language comprehension abilities.

Since no gains were observed in expressive language, we further examined vocalizations in the Study group to obtain additional insight into the intervention’s impact on expressive language.

### Vocalization analysis

Significant changes in vocalizations were observed only in the 4-to-5-year-olds group. These children spoke more, produced a greater number of content words, and used a wider variety of grammatical words across sessions (Fig. 3A), suggesting that the intervention may provide a playful context for practicing verbal behavior.

**Figure 3.**
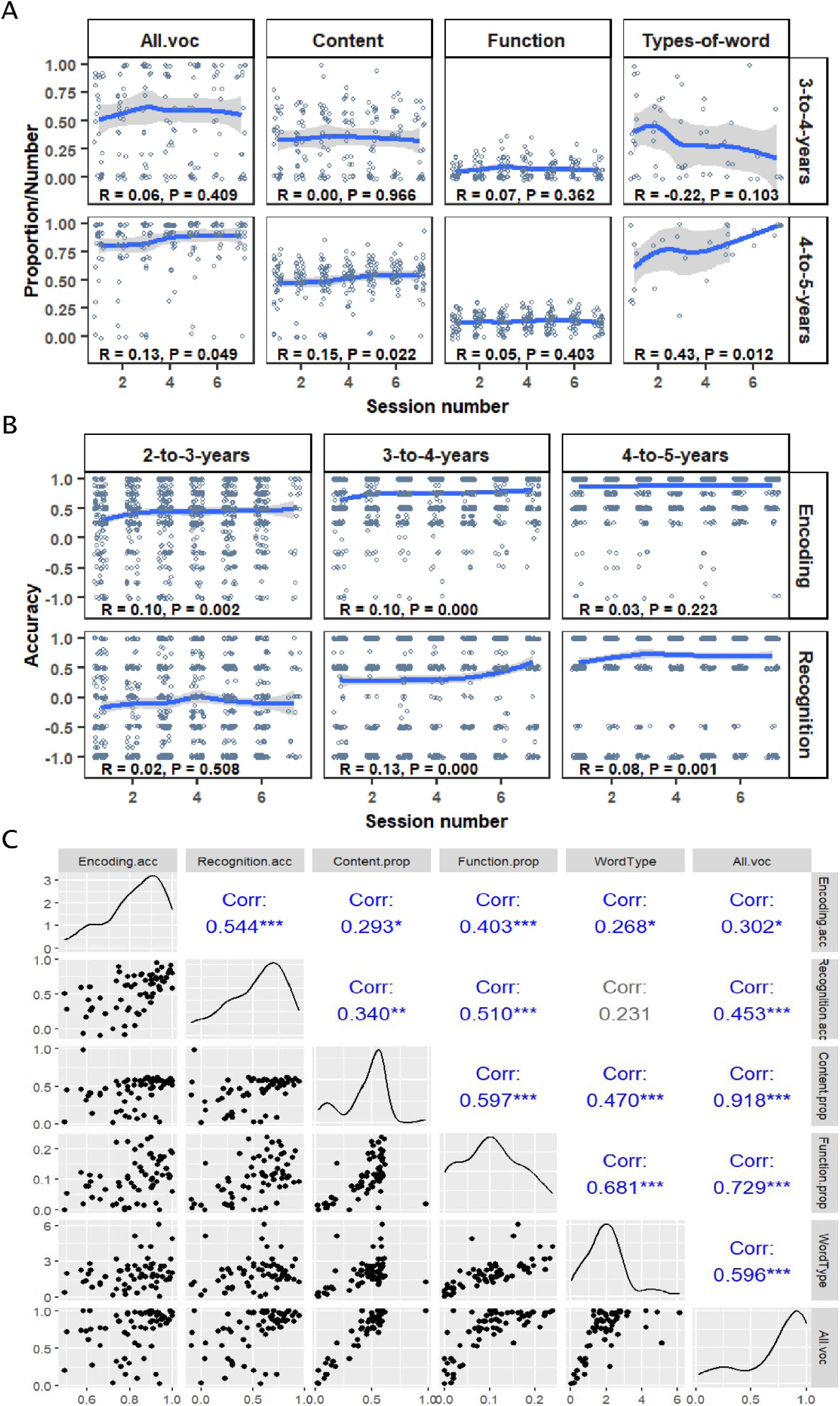
Performance during training in Study group. In **A**, we plot the proportion of word categories produced across training sessions. Panels show the changes in the proportion of total vocabulary (All.voc), content words (Content), function words (Function), and word types (Types-of-word) the child produced across sessions. Upper row: 3-4-year-olds; lower row: 4-5-year-olds. Spearman correlation coefficients (R) and corresponding *p* values indicate associations between session number and proportion of vocalized words. In **B**, we plot the accuracy in Encoding and recognition phases, obtained from tactile’s child’s responses, by age group. On the y-axis, zero corresponds to accuracy at chance. Top row: encoding accuracy; bottom row: recognition accuracy. Panels correspond to children aged 2-3 years, 3-4 years, and 4-5 years. Spearman correlation coefficients (R) and *p* values indicate associations between session number and performance accuracy. In **A** and **B**, the scatterplots display individual observations across training with locally weighted smoothing curves (blue lines) and 95% CIs (shaded areas). In **C**, we plot pairwise correlations among the performance in the encoding, recognition and vocalizations phases. Diagonal panels show variable distributions; lower panels display scatterplots; upper panels present Spearman correlation coefficients (Corr) with significance levels (* = p < .05; ** = p < .01; *** = p < .001). Variables include encoding accuracy (Encoding.acc), recognition accuracy (Recognition.acc), proportion of content words (Content.prop), proportion of function words (Function.prop), number of word types (WordType), and total vocabulary proportion (All.voc).

Additionally, across all preschoolers, the mean number of vocalizations was positively correlated with mean accuracy in both the encoding and recognition phases (Figs 3B and 3C). This suggests that children who more accurately encoded and recognized word-image associations also tended to vocalize more frequently. This relationship may reflect underlying differences in linguistic ability or task engagement, although further studies are needed to clarify the mechanisms involved. In the absence of comparable vocalization data from a control group, these findings should be interpreted as descriptive of in-task behavior rather than evidence of generalized improvements in expressive language.

### ANCOVA analysis

Finally, we examined the possibility that the observed Group differences were due to pre-existing disparities in linguistic skills by conducting nonparametric ANCOVAs. No significant differences were found in any comparison (2-3 years, Bayley language comprehension, p = 0.298; 3-4 years, TACL Morphology, p = 0.559; TACL Syntax, p = 0.202; TACL Vocabulary, p = 0.130; 4-5 years, STSG language comprehension, p = 0.312).

## Discussion

Our results indicate that a structured, tablet-based language intervention can be associated with improvements in language comprehension across both toddlerhood and preschool years. Since the groups were balanced at baseline and results remained consistent after adjustment, these differences are unlikely to stem from preexisting disparities. While concerns regarding the deleterious effects of digital media are prevalent [20], our findings indicate no such negative outcomes. On the contrary, the observation of measurable gains following an intervention of only four to eight weeks suggests helpfulness. This relatively brief and structured format is particularly well-suited for early childhood, as it prevents the cognitive overload and attentional fatigue often associated with prolonged digital exposure, thereby maximizing the quality of engagement over its quantity.

There is broad consensus that research in real-world settings using quasi-experiment designs can yield small-to-moderate effect sizes that provide meaningful insights for clinical and educational practice [21]. The interactive nature of our intervention, requiring active touchscreen input and speech production rather than passive consumption, was likely a key driver of the gains we observed. By featuring educator videos with eye contact and positive feedback, the pedagogical design mimicked the social engagement crucial for early language acquisition. This approach likely engaged multi-modal learning pathways, integrating auditory, visual, and motor inputs, fostering a contingent interaction that serves as an active learning environment rather than a mere source of digital exposure.

Overall, the intervention appears to have potentiated the natural developmental trajectory of preschoolers. During this stage, vocabulary, morphology, and syntax evolve rapidly along distinct paths. Vocabulary undergoes significant expansion not only in terms of lexical quantity [13] but also in the depth of semantic integration and the complexity of word relations [22]. As vocabulary growth is heavily contingent upon environmental quality [8,9], the tablet-based training served as a valuable enrichment. In contrast, morphosyntactic development tends to progress more gradually and exhibits less individual variability [23]. This relative stability likely stems from the fact that grammar relies on complex temporal integration and rule-based mechanisms, such as abstraction and generalization. Morphosyntax being partially mediated by vocabulary size, it is also highly sensitive to the quality of environmental stimulation [9]. Furthermore, while vocabulary and syntax are foundational to both comprehension and expression, speech production introduces the additional cognitive demand of encoding thoughts into verbal messages within socially contingent contexts. Further research with larger cohorts is required to determine whether intervention-driven gains in one area might catalyze improvements in the others.

Globally, our results indicate that effects are strongest in direct assessments, while parent-report measures may be less sensitive. The discrepancy observed between parent-reported measures (ASQ-3) and professional assessments (Bayley) is well-documented [24,25]. Even in larger cohorts, these correlations remain moderate at best, particularly when tracking nuanced longitudinal changes rather than major developmental shifts. The discrepancy might have been exacerbated in our study by the COVID-19 pandemic, where parents, sensitized by concerns over developmental delays due to lockdowns, may have adopted a more pessimistic lens [26,27]. Clinically, this divergence may also reflect a ‘lag’ in perception; receptive gains often remain ‘invisible’ to caregivers until they surface as expressive speech. Furthermore, our data confirmed that parental education was significantly associated with outcomes, particularly among older children [28-31].

This highlights the role of environmental scaffolding, where the intervention’s impact is influenced by the preexisting linguistic capital of the household.

While consistent group differences in standardized expressive measures were not observed, exploratory analyses of the Study group’s in-app vocalizations revealed that older preschoolers increased both the frequency and diversity of their speech production. Furthermore, children with higher task accuracy had greater vocalization rates. Although these behaviors did not correlate with standardized gain scores, suggesting they may reflect task-specific engagement rather than generalized linguistic advancement, the cultivation of verbal behavior remains clinically significant. Proficient spoken language is a cornerstone of social and academic success; children with robust expressive skills exhibit superior social competence, more positive peer interactions [32], and a higher propensity for prosocial behavior [33].

From a theoretical perspective, the present findings can be interpreted in light of empirically supported mechanisms of social interaction in language learning. The importance of ostensive social cues and communicative contexts in guiding early learning has been emphasized in developmental research [34]. Research in language acquisition indicates that children learn words more effectively when linguistic input occurs within episodes of shared attention, contingent responsiveness, and structured support [35]. Moreover, tasks involving immediate recognition and feedback may confer additional benefits, as previous studies have shown that preschoolers’ word learning improves with immediate retrieval within single sessions [36]. In this context, the effectiveness of the present intervention may stem not from digital exposure per se, but from its capacity to incorporate these interactional features in a playful, human-supervised environment. Thus, the intervention can be understood as a socially enriched learning environment that supports educator-child interactions rather than as a passive digital tool.

Despite these promising results, several limitations must be acknowledged. First, our assessment took place immediately post-intervention; therefore, the long-term retention of these linguistic gains remains to be determined through longitudinal follow-up. Second, while we emphasized the role of the digital tool, the socio-technical interaction between the child, the tablet, and the supervising adult likely contributed to the intervention’s success. This aligns with evidence that digital learning is most effective when the content is reinforced through real-world application during daily routines. Future research should aim to disentangle the specific contributions of adult scaffolding versus independent digital interaction to further refine and optimize educational protocols.

## Conclusions

Our study shows that a structured, tablet-based language intervention can yield small-to-moderate improvements in language comprehension across both toddlerhood and the preschool years. These gains remained robust after controlling child and parental factors, suggesting that the results are not attributable to preexisting group differences. Critically, no negative effects were detected under brief, supervised use, informing ongoing concerns about early digital media exposure.

We argue that the focus of public health and educational discourse should shift from technology itself to the modality and context of its use. When human or financial resources are limited, these devices offer a scalable and accessible alternative for developmental support. Furthermore, while further research is required to map the transition from receptive to expressive gains, our exploratory data suggest that older preschoolers may leverage the contingent nature of interactive tablets to practice verbal behavior, a skill vital for social and academic success. Ultimately, tablet-based tools, when pedagogically structured and supervised, represent a valuable complement to naturalistic language development rather than a replacement for it.

## Supporting information

Table presenting the statistical comparison between children included in the analysis and those excluded due to missing post-training evaluation data

## Acknowledgments

We are grateful to the families, the Servicio de Salud Metropolitano Sur Oriente. M.P. received support from the grants FONDECYT #1241946, FONDEF IT24I0139, and National Center for Artificial Intelligence CENIA FB210017, Basal ANID. G.D.L. was supported by the French government managed by the Agence Nationale de la Recherche as part of the France 2030 program (ANR-23-IAHU-0010). P.G. was partially supported by ANID AC3E CIA250006. The authors used an AI-assisted language tool to improve the clarity and grammar of the manuscript. All scientific content and interpretations are the sole responsibility of the authors.

## Supporting information

**Table S1**.

