## Supplementary material for "A supervised digital game intervention supports language and communication in young children": Table presenting the statistical comparison between children included in the analysis and those excluded due to missing post-training evaluation data

**Supporting information**

**Table S1. Comparison of the children included and excluded in the analysis.** The included children provided at least one pre-post evaluation in one or more assessment. Excluded children had only one pre-training assessment. ^a^: we did not report the p value when n was less than seven.

| **(A) Demographic data** | | | | | | | | | | | | | | | | |
| --- | --- | --- | --- | --- | --- | --- | --- | --- | --- | --- | --- | --- | --- | --- | --- | --- |
|  | | | **2-to-3** | | | |  | **3-to-4** | | | |  | **4a5** | | | |
|  |  |  | **n** | **Mean** | **SD** | ***p*** |  | **n** | **Mean** | **SD** | ***p*** |  | **n** | **Mean** | **SD** | ***p*** |
| Child age | Control | Included | 41 | 29.26 | 3.50 | 0.413 |  | 49 | 41.46 | 3.80 | 0.075 |  | 37 | 55.73 | 5.51 | 0.416 |
|  |  | Excluded | 17 | 30.29 | 4.56 |  |  | 19 | 43.12 | 3.19 |  |  | 9 | 53.74 | 6.51 |  |
|  | Study | Included | 47 | 28.04 | 4.05 | 0.083 |  | 38 | 42.72 | 3.24 | 0.584 |  | 37 | 55.92 | 5.45 | a |
|  |  | Excluded | 16 | 29.89 | 3.39 |  |  | 17 | 43.17 | 2.57 |  |  | 4 | 60.14 | 3.27 |  |
| Mother age | Control | Included | 41 | 30.24 | 5.77 | 0.601 |  | 41 | 32.73 | 5.78 | 0.092 |  | 37 | 34.03 | 5.88 | 0.244 |
|  |  | Excluded | 16 | 31.13 | 5.60 |  |  | 16 | 30.53 | 4.22 |  |  | 9 | 31.44 | 5.64 |  |
|  | Study | Included | 47 | 31.53 | 6.11 | 0.070 |  | 47 | 31.84 | 6.57 | 0.532 |  | 37 | 33.46 | 6.19 | ^a^ |
|  |  | Excluded | 13 | 34.62 | 4.91 |  |  | 13 | 33.13 | 6.65 |  |  | 4 | 83.75 | 98.99 |  |
| Mother education | Control | Included | 39 | 5.69 | 1.92 | 0.295 |  | 48 | 5.38 | 1.86 | 0.814 |  | 36 | 6.17 | 1.70 | 0.930 |
|  |  | Excluded | 16 | 6.25 | 1.69 |  |  | 18 | 5.50 | 1.92 |  |  | 9 | 6.22 | 1.64 |  |
|  | Study | Included | 47 | 5.74 | 1.89 | 0.261 |  | 33 | 5.33 | 1.74 | 0.638 |  | 37 | 5.43 | 1.83 | ^a^ |
|  |  | Excluded | 12 | 5.00 | 2.00 |  |  | 16 | 5.63 | 2.13 |  |  | 4 | 7.00 | 1.15 |  |
| Father age | Control | Included | 39 | 32.08 | 6.74 | 0.967 |  | 48 | 33.63 | 6.23 | 0.562 |  | 36 | 36.22 | 6.73 | 0.111 |
|  |  | Excluded | 16 | 32.00 | 6.11 |  |  | 18 | 32.50 | 7.17 |  |  | 9 | 32.44 | 5.75 |  |
|  | Study | Included | 47 | 33.90 | 6.55 | 0.383 |  | 33 | 33.21 | 7.56 | 0.211 |  | 37 | 36.41 | 7.68 | ^a^ |
|  |  | Excluded | 12 | 36.08 | 7.96 |  |  | 16 | 36.33 | 7.93 |  |  | 4 | 35.75 | 10.69 |  |
| Father education | Control | Included | 36 | 5.78 | 1.84 | 0.617 |  | 46 | 5.04 | 1.93 | 0.486 |  | 34 | 5.53 | 1.94 | 0.923 |
|  |  | Excluded | 16 | 5.50 | 1.83 |  |  | 17 | 5.41 | 1.80 |  |  | 8 | 5.63 | 2.56 |  |
|  | Study | Included | 43 | 5.47 | 1.87 | 0.431 |  | 32 | 5.47 | 1.87 | 0.202 |  | 37 | 5.30 | 2.22 | ^a^ |
|  |  | Excluded | 13 | 6.00 | 2.16 |  |  | 16 | 4.63 | 2.22 |  |  | 4 | 6.50 | 1.00 |  |
| **(B) ASQ-3** | | | | | | | | | | | | | | | | |
|  | | | **2-to-3** | | | |  | **3-to-4** | | | |  | **4a5** | | | |
|  |  |  | **n** | **Mean** | **SD** | ***p*** |  | **n** | **Mean** | **SD** | ***p*** |  | **n** | **Mean** | **SD** | ***p*** |
| ASQ-3 Com | Control | Included | 41 | 41.95 | 15.28 | 0.230 |  | 48 | 50.83 | 12.90 | 0.277 |  | 37 | 52.57 | 11.52 | 0.216 |
|  |  | Excluded | 15 | 36.00 | 16.28 |  |  | 16 | 45.63 | 17.11 |  |  | 9 | 47.22 | 10.93 |  |
|  | Study | Included | 47 | 45.21 | 13.79 | 0.795 |  | 36 | 46.25 | 13.44 | 0.131 |  | 37 | 52.43 | 9.47 | ^a^ |
|  |  | Excluded | 13 | 44.23 | 11.34 |  |  | 16 | 51.56 | 10.44 |  |  | 4 | 53.75 | 7.50 |  |
| ASQ-3 GM | Control | Included | 41 | 48.78 | 11.71 | 0.762 |  | 48 | 52.08 | 8.74 | 0.540 |  | 37 | 53.78 | 8.69 | 0.560 |
|  |  | Excluded | 15 | 49.67 | 8.76 |  |  | 17 | 50.59 | 8.46 |  |  | 9 | 51.67 | 9.68 |  |
|  | Study | Included | 46 | 52.17 | 9.05 | 0.701 |  | 36 | 50.69 | 8.88 | 0.775 |  | 37 | 54.05 | 8.07 | ^a^ |
|  |  | Excluded | 13 | 53.46 | 10.88 |  |  | 16 | 51.56 | 10.44 |  |  | 4 | 43.75 | 17.02 |  |
| ASQ-3 FM | Control | Included | 41 | 36.22 | 13.77 | 0.222 |  | 48 | 39.69 | 15.38 | 0.056 |  | 37 | 46.22 | 13.04 | 0.607 |
|  |  | Excluded | 15 | 41.00 | 12.28 |  |  | 17 | 30.59 | 16.38 |  |  | 9 | 43.33 | 15.00 |  |
|  | Study | Included | 46 | 40.11 | 14.12 | 0.202 |  | 36 | 35.42 | 17.86 | 0.552 |  | 37 | 43.78 | 16.09 | ^a^ |
|  |  | Excluded | 13 | 33.85 | 15.30 |  |  | 16 | 38.44 | 16.20 |  |  | 4 | 53.75 | 9.46 |  |
| ASQ-3 PrS | Control | Included | 41 | 40.98 | 14.88 | 0.367 |  | 48 | 50.83 | 9.80 | 0.063 |  | 37 | 52.16 | 9.83 | 0.984 |
|  |  | Excluded | 15 | 44.67 | 12.74 |  |  | 16 | 43.13 | 14.59 |  |  | 9 | 52.22 | 7.55 |  |
|  | Study | Included | 46 | 44.89 | 12.09 | 0.276 |  | 36 | 48.33 | 12.48 | 0.953 |  | 37 | 48.92 | 14.10 | ^a^ |
|  |  | Excluded | 13 | 48.85 | 11.02 |  |  | 16 | 48.13 | 11.38 |  |  | 4 | 55.00 | 4.08 |  |
| ASQ-3 PeS | Control | Included | 41 | 41.34 | 12.85 | 0.523 |  | 48 | 48.13 | 11.74 | 0.171 |  | 37 | 52.03 | 8.37 | 0.236 |
|  |  | Excluded | 15 | 39.00 | 11.68 |  |  | 17 | 42.35 | 15.32 |  |  | 9 | 48.33 | 7.91 |  |
|  | Study | Included | 46 | 44.35 | 10.31 | 0.211 |  | 36 | 48.33 | 9.93 | 0.587 |  | 37 | 53.51 | 7.44 | ^a^ |
|  |  | Excluded | 13 | 49.23 | 12.39 |  |  | 16 | 50.00 | 10.17 |  |  | 4 | 45.00 | 7.07 |  |
| **(C) Linguistic abilities** | | | | | | | | | | | | | | | | |
|  | | | **2-to-3** | | | |  | **3-to-4** | | | |  | **4a5** | | | |
|  |  |  | **n** | **Mean** | **SD** | ***p*** |  | **n** | **Mean** | **SD** | ***p*** |  | **n** | **Mean** | **SD** | ***p*** |
| Bayley C | Control | Included | 36 | 28.67 | 5.82 | 0.900 |  |  |  |  |  |  |  |  |  |  |
|  |  | Excluded | 4 | 27.75 | 13.30 |  |  |  |  |  |  |  |  |  |  |  |
|  | Study | Included | 33 | 27.03 | 5.85 | 0.344 |  |  |  |  |  |  |  |  |  |  |
|  |  | Excluded | 11 | 24.64 | 7.38 |  |  |  |  |  |  |  |  |  |  |  |
| Bayley E | Control | Included | 36 | 26.19 | 7.71 | 0.913 |  |  |  |  |  |  |  |  |  |  |
|  |  | Excluded | 4 | 25.25 | 15.84 |  |  |  |  |  |  |  |  |  |  |  |
|  | Study | Included | 33 | 25.52 | 6.75 | 0.153 |  |  |  |  |  |  |  |  |  |  |
|  |  | Excluded | 11 | 21.91 | 6.98 |  |  |  |  |  |  |  |  |  |  |  |
| STSG_E | Control | Included |  |  |  |  |  | 35 | 11.46 | 8.99 | 0.817 |  | 33 | 24.33 | 10.53 | ^a^ |
|  |  | Excluded |  |  |  |  |  | 6 | 9.83 | 15.96 |  |  | 2 | 2.50 | 2.12 |  |
|  | Study | Included |  |  |  |  |  | 28 | 10.86 | 10.37 | ^a^ |  | 33 | 21.73 | 10.91 | ^a^ |
|  |  | Excluded |  |  |  |  |  | 3 | 14.67 | 9.29 |  |  | 2 | 18.00 | 9.90 |  |
| STSG_C | Control | Included |  |  |  |  |  | 35 | 23.46 | 8.31 | ^a^ |  | 33 | 28.88 | 8.31 | ^a^ |
|  |  | Excluded |  |  |  |  |  | 5 | 16.80 | 14.60 |  |  | 2 | 23.00 | 14.60 |  |
|  | Study | Included |  |  |  |  |  | 28 | 20.79 | 9.93 | ^a^ |  | 33 | 28.45 | 9.93 | ^a^ |
|  |  | Excluded |  |  |  |  |  | 3 | 23.67 | 4.51 |  |  | 2 | 30.50 | 4.51 |  |
| Tacl_t | Control | Included |  |  |  |  |  | 28 | 59.14 | 16.86 | ^a^ |  |  |  |  |  |
|  |  | Excluded |  |  |  |  |  | 2 | 66.50 | 9.19 |  |  |  |  |  |  |
|  | Study | Included |  |  |  |  |  | 22 | 52.77 | 13.05 | ^a^ |  |  |  |  |  |
|  |  | Excluded |  |  |  |  |  | 3 | 60.67 | 11.72 |  |  |  |  |  |  |
| Tacl_V | Control | Included |  |  |  |  |  | 28 | 27.57 | 7.25 | ^a^ |  |  |  |  |  |
|  |  | Excluded |  |  |  |  |  | 2 | 30.00 | 4.24 |  |  |  |  |  |  |
|  | Study | Included |  |  |  |  |  | 22 | 25.41 | 5.15 | ^a^ |  |  |  |  |  |
|  |  | Excluded |  |  |  |  |  | 3 | 28.67 | 1.53 |  |  |  |  |  |  |
| Tacl_M | Control | Included |  |  |  |  |  | 28 | 25.96 | 8.83 | ^a^ |  |  |  |  |  |
|  |  | Excluded |  |  |  |  |  | 2 | 30.00 | 4.24 |  |  |  |  |  |  |
|  | Study | Included |  |  |  |  |  | 22 | 22.68 | 8.31 | ^a^ |  |  |  |  |  |
|  |  | Excluded |  |  |  |  |  | 3 | 26.67 | 8.33 |  |  |  |  |  |  |
| Tacl_S | Control | Included |  |  |  |  |  | 28 | 5.61 | 2.17 | ^a^ |  |  |  |  |  |
|  |  | Excluded |  |  |  |  |  | 2 | 6.50 | 0.71 |  |  |  |  |  |  |
|  | Study | Included |  |  |  |  |  | 22 | 4.68 | 2.32 | ^a^ |  |  |  |  |  |
|  |  | Excluded |  |  |  |  |  | 3 | 5.33 | 2.52 |  |  |  |  |  |  |
